# Exploring the Residue-Level Interactions between the R2ab Protein and Polystyrene Nanoparticles

**DOI:** 10.1101/2023.08.28.554951

**Authors:** Radha P. Somarathne, Sandeep K. Misra, Chathuri S. Kariyawasam, Jacques J. Kessl, Joshua S. Sharp, Nicholas C. Fitzkee

**Affiliations:** Department of Chemistry, Mississippi State University, Mississippi State, MS 39762; Department of BioMolecular Sciences, University of Mississippi, University, MS 38677; Department of Chemistry and Biochemistry, University of Southern Mississippi, Hattiesburg, MS 39406; Department of Chemistry and Biochemistry, University of Mississippi, University, MS 38677

**Keywords:** nanoparticle interaction corona adsorbotope protein structure

## Abstract

In biological systems, proteins can bind to nanoparticles to form a “corona” of adsorbed molecules. The nanoparticle corona is of high interest because it impacts the organism’s response to the nanomaterial. Understanding the corona requires knowledge of protein structure, orientation, and dynamics at the surface. Ultimately, a residue-level mapping of protein behavior on nanoparticle surfaces is needed, but this mapping is difficult to obtain with traditional approaches. Here, we have investigated the interaction between R2ab and polystyrene nanoparticles (PSNPs) at the level of individual residues. R2ab is a bacterial surface protein from *Staphylococcus epidermidis* and is known to interact strongly with polystyrene, leading to biofilm formation. We have used mass spectrometry after lysine methylation and hydrogen-deuterium exchange (HDX) NMR spectroscopy to understand how the R2ab protein interacts with PSNPs of different sizes. Through lysine methylation, we observe subtle but statistically significant changes in methylation patterns in the presence of PSNPs, indicating altered protein surface accessibility. HDX measurements reveal that certain regions of the R2ab protein undergo faster exchange rates in the presence of PSNPs, suggesting conformational changes upon binding. Both results support a recently proposed “adsorbotope” model, wherein adsorbed proteins consist of unfolded anchor points interspersed with regions of partial structure. Our data also highlight the challenges of characterizing complex protein-nanoparticle interactions using these techniques, such as fast exchange rates. While providing insights into how proteins respond to nanoparticle surfaces, this research emphasizes the need for advanced methods to comprehend these intricate interactions fully at the residue level.

**TOC Image:** 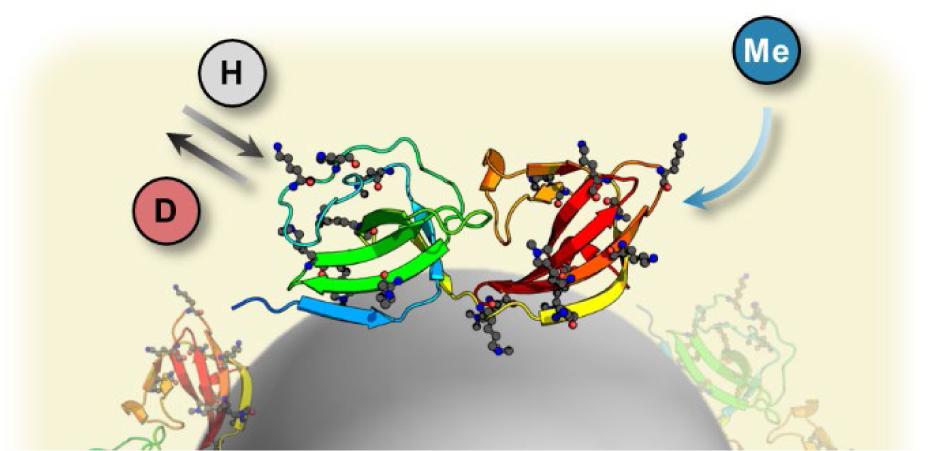

Lysine methylation and hydrogen-deuterium exchange can reveal useful structural details about protein adsorption to nanoparticle surfaces.

## Introduction

The nanoparticle protein corona remains a major challenge to developing a predictive understanding of bio-nano interactions. In biological fluids, the proteins that spontaneously adsorb to nanomaterials, can either bind loosely (“soft” corona) or tightly (“hard” corona), and the resulting corona influences how the organism responds.^1–4^ Polystyrene nanoparticles (PSNPs), while limited in their applications as a drug delivery tool, serve as a useful model for understanding the nanoparticle corona.^5–7^ However, understanding the interaction of proteins with PSNPs requires investigating the protein structure, orientation, and dynamics. Techniques such as circular dichroism and calorimetric studies have been invaluable,^5,8–10^ but developing a residue-based understanding of adsorption remains extremely challenging. Alternative strategies are needed to determine which particular regions of a protein may be perturbed upon interaction with PSNPs.

One potential tool for studying how proteins are perturbed in the presence of PSNPs is lysine methylation. Methylation is a crucial post-translational modification that is important in regulating diverse cellular processes and orchestrating complex biological functions.^11,12^ Among its various substrates, lysine residues stand out as prime targets for methylation, a chemical alteration that assigns proteins with new structural conformations and functional properties. Protein methylation of lysines contributes to gene expression regulation, signal transduction, chromatin organization, and a spectrum of cellular pathways.^13,14^ Methylation involves the transfer of a methyl group to specific lysine residues, catalyzed by a family of enzymes known as lysine methyltransferases.^15^ Methylation can also be performed reductively using formaldehyde under mild conditions that do not alter the protein’s structure.^16^ This alteration causes a distinct change in the mass of the protein, like a unique signature that can be detected through mass spectrometry.^17^ Upon proteolysis, the change in peptide mass can be used to identify the location where methylation occurred.

Understanding the pattern of methylation on a protein can provide clues about its spatial arrangement. For instance, methylation sites can reveal how the protein interacts with other molecules like DNA, RNA, or other proteins.^18,19^ If certain spots on the protein are methylated, it suggests they’re available for interactions, potentially on the protein’s surface, while unmethylated spots might be sterically occluded. The methylation pattern can also offer insight into higher-level protein structures.^20^ In proteins with multiple flexible domains, the pattern of methylation can indicate the presence of domain motions. In principle, a large, folded protein stably adsorbed to a nanoparticle surface would show a different methylation pattern than a protein in solution: The PSNP would limit methylation on the surface of contact.

Another attractive approach for monitoring protein structure in the presence of PSNPs is hydrogen-deuterium exchange (HDX).^21,22^ HDX rates can reveal which regions of a protein are exposed to solvent or possess low local stability,^23–26^ providing information that complements lysine methylation. The basic principle of HDX involves exposing a protein to a solution containing deuterium oxide (D_2_O) for a certain period of time. Deuterium atoms gradually replace labile hydrogen atoms in the protein’s backbone amide groups and side chains. The exchange rate of hydrogen and deuterium is influenced by factors such as solvent accessibility and hydrogen bonding, which in turn reflect the protein’s structural and dynamic properties. NMR can measure rapid HDX, and it is an attractive tool for characterizing proteins in the presence of nanoparticles.^27^ A protein-nanoparticle mixture is rapidly diluted in D_2_O, and real-time HDX is measured using fast-pulsing experiments like SOFAST-HMQC.^28,29^ Exchange is measured in the absence of nanoparticles, and the resulting rates are compared. No difference is expected when only a hard corona is present because adsorbed proteins do not desorb; only the non-exchanging solution rates are detected.^28^ However, if a soft corona is present, and a significant amount of proteins undergo exchange, a difference in HDX rates should be detectable.

By comparing HDX under various conditions, it is possible to create a map of deuterium incorporation, which provides information about the protein’s solvent accessibility, flexibility, and stability at different regions.^30,31^ This information is valuable for determining local and global protein folding, conformational changes, and interaction interfaces. When proteins interact with ligands and other small molecules such as nanoparticles, any conformational dynamics of proteins can also be detected, and this too could cause changes in hydrogen/deuterium exchange due to local decreases in amide exchanges, arising from decreased solvent accessibility of surface amides.^29,32,33^ The phenomenon of reduced solvent accessibility caused by ligand binding is very informative and reliable, and therefore this principle is now even used in high-throughput drug screening.^34^ As with methylation, there are obvious potential advantages when measuring HDX rates in the presence and absence of nanoparticles, and such information could potentially generate the residue-specific information needed to characterize surface-bound proteins.

In this study, we explore the residue-specific interactions of the R2ab protein in the PSNP protein corona. R2ab is a domain from the autolysin protein of *S. epidermidis* and is known to facilitate biofilm formation on polystyrene surfaces.^35–37^ R2ab is also known to interact strongly with PSNPs.^5,8^ We used quantitative measures of lysine methylation, detected by liquid chromatography-mass spectrometry (LC-MS/MS) to examine changes in methylation patterns in the presence of PSNPs. Then, we compared HDX rates in the presence and absence of PSNPs to monitor R2ab behavior. Several intriguing trends were observed, including behavior that was consistent with substantial unfolding on the PSNP surface. This work demonstrates the value of residue-based approaches for investigating the protein corona, and it highlights the particular challenges and limitations that are associated with the measurement of methylation and HDX behavior for proteins in the soft corona.

## Experimental

### Protein purification and nanoparticle characterization

Expression and purification of R2ab, along with the characterization of PSNPs, were performed as described previously.^5^ Non-functionalized polystyrene nanoparticles with varying diameters of 50 nm, 100 nm, and 200 nm were purchased from PolySciences (#08691-10, #00876-15, and #07304-15, respectively). Nanoparticle preparation was performed by dialyzing in 1L of milli-Q water, involving three exchanges where each exchange occurred for at least four hours each time.

### Methylation and LC-MS/MS analysis

R2ab was added in excess to the polystyrene nanoparticles of varying diameters with an initial concentration of 100 µM. The concentration of nanoparticles was varied to keep the total surface area constant. The PSNP concentrations used were 88, 22, and 6 nM for 50, 100, and 200 nm PSNPs, respectively. R2ab was allowed to interact with the nanoparticles for 2 hours. The samples were then centrifuged for 10 mins at 15,000 g. The supernatant was discarded, and the remaining pellet was washed three times with 20 mM Na_3_PO_4_ and 50 mM NaCl. The methylation kit was purchased from Hampton Research. The pellet was resuspended in 1 mL of buffer, and the methylation reaction of R2ab was performed according to the manufacturer’s protocol. Briefly, to each sample, 40 µL of 1M Dimethylamine Borane Complex and 80 µL of formaldehyde were added. The reaction was allowed to proceed, and after 10 minutes, the reactions were quenched with 125 µL of 1M glycine. The samples were then centrifuged for 10 mins at 15,000 g and resuspended in 100 µL of buffer. The final R2ab concentration was estimated to be 50 µM using the BCA assay (Pierce #23225). All methylation reactions were performed in triplicate.

The mass spectrometry data collection and analysis were performed in collaboration with the Glycoscience Center of Research Excellence (GlyCORE) at the University of Mississippi. The protein samples were diluted into 100 mM Tris containing 10 mM CaCl_2_ (pH 8.0). Samples were heated at 95 °C for 20 min and then placed immediately on ice for 2 min. For protein digestion, a 1:20 ratio of chymotrypsin/R2ab was added to the samples and incubated at 37 °C overnight with rotation. Chymotrypsin was inactivated by heating samples at 95 °C for 10 min and 0.1% formic acid was added to the samples before LC−MS/MS analysis. The peptides were analyzed by an Orbitrap Exploris 240 mass spectrometer coupled with a Dionex Ultimate 3000 system nanoLC system (Thermo Fisher, CA). The peptides were first loaded onto a trap column (Acclaim PepMap C18 5 µm, 0.3 × 5 mm) and then eluted onto a nano C18 column (Acclaim 3 µm, 0.075 × 150 mm) at a flow rate of 0.3 µL/min. Solvent A was 0.1% formic acid in LC-MS grade water, and solvent B was 0.1% formic acid in LC-MS grade acetonitrile. The gradient consisted of starting with 2% solvent B for 2 min, 2−32% solvent B over 13 min, ramped to 95% solvent B over 2 min and held for 2 min, returned to 2% solvent B over 2 min and held for 7 min. Electrospray ionization was carried out in a positive ion mode, with a nominal orbitrap resolution of 60,000. Data-dependent mode MS/MS was used, with an HCD energy of 35, and the top 8 peaks were selected for fragmentation. Byonic (v4.4.1, Protein Metrics, San Carlos, CA) was used to identify the peptides and determine the protein sequence coverage. The extent of methylation on lysine residues per peptide was calculated manually with the help of Xcalibur V.4.5 by dividing the area under the curve of methylated versions of that peptide with the sum of methylated and unmethylated area under the curve of that peptide. The mass spectrometry proteomics data have been deposited to the ProteomeXchange Consortium via the PRIDE partner repository with the dataset identifier PXD044469.^38^

### Sample preparation for NMR

NMR samples were prepared with 200 µM R2ab and 50 µM of 50 nm non-functionalized PSNPs in 50 mM sodium acetate buffer at pH 4.5, along with 18.2 MΩ cm ultrapure Milli-Q water, to bring the final volume to 550 µL. Assignments were transferred from spectra collected at pH 6.5 (BMRB ID 30821). The spectrum R2ab at pH 4.5 yielded 43 well-defined and transferable assignments (Supporting Information, **Figure S1**). Then, the protein/PSNP mixture was transferred to a clean NMR tube (Wilmad, #535-PP-7).

### Measurement of hydrogen-deuterium exchange (HDX) NMR

All NMR experiments were performed on a 600 MHz Bruker Avance III NMR system equipped with a CP-QCI cryoprobe and experiments were recorded at 25 °C. A sensitivity-enhanced SOFAST-HMQC spectrum was taken after tuning and 3D shimming,^29,39,40^ and the peak intensities of this control sample were recorded. Then, to this sample, 550 µL of D_2_0, was added to the sample, and the sample was immediately vortexed, and the NMR tube was loaded into the NMR. Immediately after a quick 1D shimming, a series of SOFAST-HMQC spectra were continuously taken to monitor the change in peak intensities during binding. Because of the dilution, the final protein concentration was approximately 100 μM. The dead time (from the addition of D_2_O to the onset of data acquisition) was less than five minutes. Note that, in this experiment the final concentration of D_2_O is approximately 50%, meaning that peaks should decay to a baseline of approximately 25% of their original intensity (following a 1:2 dilution and exchange with 50% D_2_O). This information was used in the final fitting, where peaks were fit to the equation:

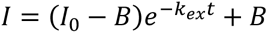

In this equation, *I* is the observed intensity as a function of time, *I*_0_ represents the initial intensity which is constrained during the fit; *I*_0_ is estimated from the initial intensity. The apparent exchange rate *k*_*ex*_ and the baseline *B* were determined by nonlinear least squares fitting using Gnuplot. Uncertainties were estimated from the NMR spectral noise and calculated using Gnuplot.

For this SOFAST-HMQC experiment the indirect acquisition time (t1) was 120 ms, recorded for a total of 64 complex points. The direct acquisition time (t2) was 120 ms. A total of 16 scans were performed, with each experiment lasting for approximately 5 minutes. All spectral data were processed using NMRPipe and POKY. All processed spectra, along with fitting scripts and analysis output, have been deposited in Zenodo (https://doi.org/10.5281/zenodo.8284557).

## Results & Discussion

### Methylation of R2ab in the Presence of PSNPs Shows Several Distinct Classes of Accessibility

Methylation of proteins plays a pivotal role in various cellular processes, and its detection and analysis have become essential in the field of proteomics.^41^ As a chemical modification, methylation can also be helpful in characterizing transient protein states.^42^ Previously, we found that lysine methylation could be used to modulate binding to gold nanospheres.^43^ We started our study by hypothesizing that lysine methylation could be used to determine protein orientation and structure on a nanoparticle.^44^ Lysine, with its lone-pair electrons and its frequent presence on the protein surface, is particularly prone to a variety of posttranslational modifications.^45,46^ Lysine methylation stands out from other lysine modifications for two primary reasons: First, each methyl group addition retains the +1 charge of the amine at physiological pH, unlike other modifications that transform the charged amine into a neutral amide.^47^ Second, among all posttranslational modifications, lysine methylation has the least impact on the side chain’s overall size.^48^ These two factors mean that methylation of lysine residues is a fairly mild chemical alteration that is still detectable using mass-spectrometry-based proteomics approaches.

R2ab is a dynamic protein consisting of two main subdomains, R2a and R2b. It contains an abundance of lysines, which are evenly distributed throughout its structure. Some of these lysine residues are exposed to the surrounding solution, while others remain partially buried. In the presence of PSNPs, R2ab has been observed to undergo some unfolding, indicating its sensitivity to the nanoparticle environment. To study the effects of PSNPs on R2ab, we methylated the protein in its native form and exposed it to PSNPs of various diameters. The exposure time of for methylation was brief (5 min), allowing a comparison between the PSNP-exposed R2ab and the R2ab domain in the absence of PSNPs. This work is similar to what was done by McClain, *et al.*, who used proteolysis to monitor structural changes occurring during protein-nanoparticle interactions.^49^

Methylation, followed by proteolytic digestion, identified a total of 9 major fragments in R2ab (**Figure 1**). By quantitatively integrating each fragment signal in the HPLC chromatogram, we could compare methylation rates in the presence and absence of PSNPs. Overall, the degree of methylation is high, although small, statistically different values are observed. The small differences in methylation levels in the presence of PSNPs are likely due to the protein’s ability to quickly respond to the nanoparticle environment and the partial unfolding that occurs on the nanoparticle surface.^8^ All peptide fragments identified by LC-MS contain one or more lysine residues (**Figure 1**). While using HPLC-based quantitation allows accurate comparison of each peptide, we were not able to exhaustively identify every lysine in R2ab. In addition, varying degrees of methylation make it impossible to determine the methylation state for every lysine in every fragment. To address this limitation in our study, we assumed that the impact of multiple lysines within a given fragment is similar. This assumption enables us to make meaningful comparisons despite the complexity arising from multiple lysine residues in each peptide.

**Figure 1.**
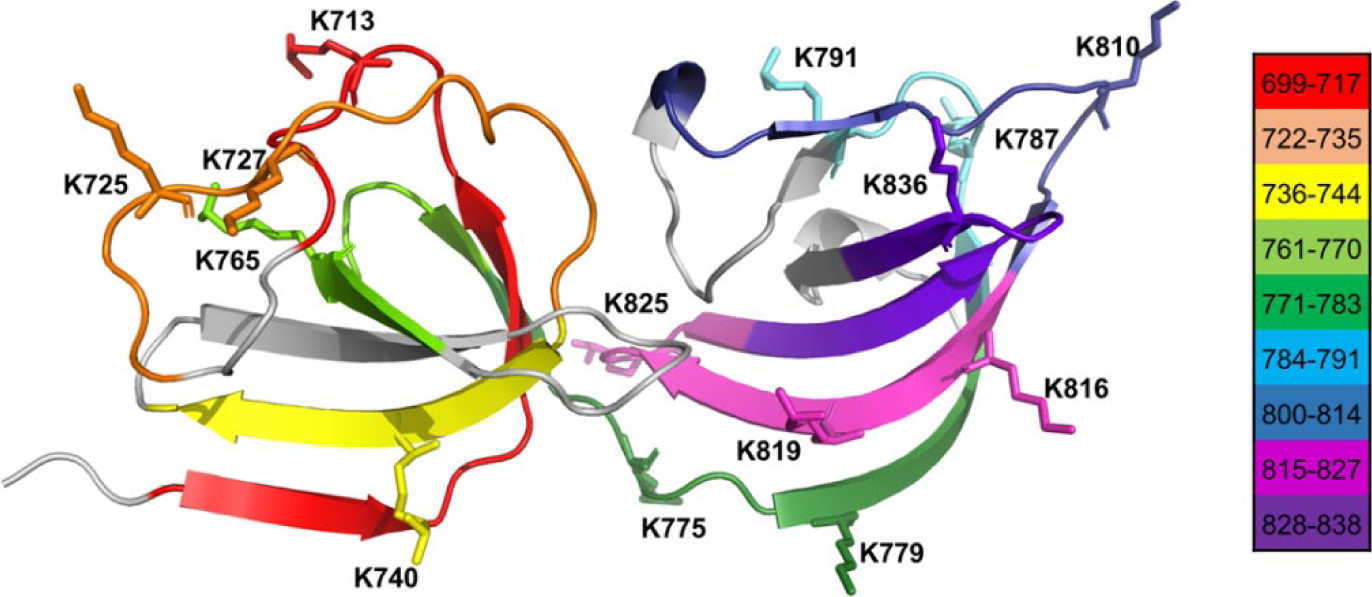
Peptide fragments of R2ab and the lysines in the respective fragments that are detected by mass spectrometry are represented by unique colors for identification.

It is possible to categorize peptide fragments of R2ab based on the change in their methylation levels in the presence of PSNPs (**Figure 2A**). Among the fragments analyzed, we found that fragments 699-717, 736-744, 771-783, 800-814, and 828-838 display lower degrees of methylation when exposed to PSNPs. In contrast, fragment 722-735 shows a higher degree of methylation in the presence of PSNPs. Meanwhile, fragments 761-770, 784-791, and 815-827 do not appear to be affected by the presence of PSNPs. The observed lower degrees of methylation in certain peptide fragments suggest that these regions are likely less accessible following the interaction with the PSNPs. For example, fragment 800-814, located in the R2b subdomain, contains one lysine (K810), and in the absence of PSNPs state, it is entirely methylated. However, the presence of PSNPs leads to a small but statistically significant decrease in methylation, indicating that this region of the R2b subdomain interacts more strongly with the PSNPs. However, this reduced methylation does not seem to be influenced by the curvature of the nanoparticles (**Figures 2B-E**), since there is no trend seen for these residues as the diameter is increased from 50-200 nm. On the other hand, fragment 784-791 in the R2b chain is easily accessible from the rear face of the protein and is fully methylated in the absence of PSNPs. The presence of nanoparticles does not appear to alter its methylation level, suggesting that this region remains accessible when adsorbed to PSNP surfaces. Peptide fragment 761-770, primarily located in the R2a chain, acts as the connecting link between the two chains. Although it contains one lysine in a loop, the entire fragment is sandwiched between the two chains, making it less accessible to methylation, regardless of the presence of nanoparticles. Consequently, the overall degree of methylation for this fragment remains lower than the others.

**Figure 2.**
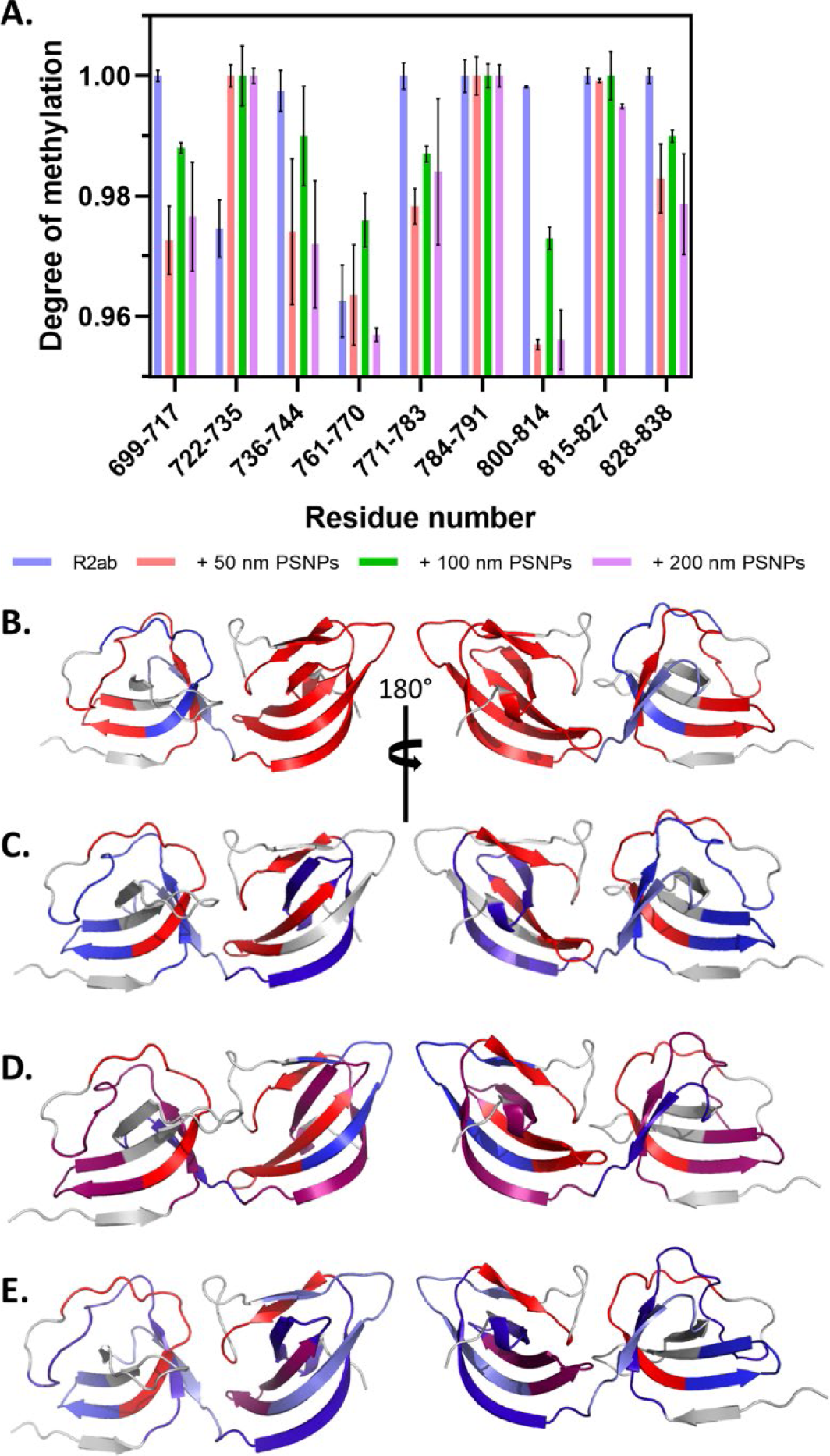
Degrees of methylation show only subtle changes regardless of the presence of PSNPs **(A)**. Error bars represent the standard deviation from three separate experiments. For comparison, regions of R2ab have been colored to show how well the individual peptide fragments have been methylated in the native form **(B)**, in the presence of 50 nm **(C)**, 100 nm **(D)** and 200 nm **(E)** PSNPs. Regions which are change from being more methylated to less methylated, (red to blue) indicate possible changes in protein orientation and disorder.

Finally, we observe that some fragments are ambiguous. Such an example is fragment 736-744, which appears to exhibit lower methylation in the presence of PSNPs but exhibits a large deviation between the three measurements. All of the fragments had a high degree of methylation, and because of the time needed to mix R2ab with PSNPs, perform the methylation, displace the proteins, and perform a proteolytic digest, shorter times were not feasible. Thus, while the observed differences in methylation are reproducible, they are also small. This represents one limitation of this approach; faster labeling methods, such as fast photochemical oxidation of proteins (FPOP),^50,51^ or using proteolysis alone,^49^ may be better suited for probing the orientation and accessibility of proteins interacting with nanoparticles.

### Structural Interpretation of Methylated R2ab in the Presence of PSNPs

It is tempting to interpret our results using a structural approach; indeed, if R2ab remains folded when near the PSNP surface, we expect that lysine residues will experience steric occlusion and a lower degree of methylation. This is complicated by two factors: First, it is known that R2ab samples an equilibrium between the adsorbed-desorbed states on PSNPs,^52–54^ (and this exchange likely occurs on the order of seconds or faster. Thus, the observed changes in the degree of methylation will reflect a weighted average of the bound vs. the unbound state. Second, it is likely that proteins partially unfold upon contact with a PSNP surface.^8^ Previously, we used the term “adsorbotope” to describe a peptide region that has a higher-than-normal affinity for a nanoparticle surface. These high-affinity adsorbotopes are interspersed with regions of the non-interacting sequence, which could be ordered or disordered. On PSNPs, we found strong evidence that much of the protein appeared to be disordered.

Thus, while fragment 800-814 (**Figure 2A**) may represent a protein conformation where a folded R2b subdomain is occluded by the nanoparticle surface, it is equally possible that this fragment contains an adsorbotope site and the surrounding sequence is disordered. Similarly, while fragment 784-791 may exist in an orientation where this region is fully exposed to solvent in the presence of PSNPs, it is equally likely that this region unfolds in the presence of PSNPs and retains its accessibility, despite the fact that the next region (800-814) is more occluded. Using this methylation data alone, it is impossible to distinguish these two possibilities. At best, we can say that fragment 800-814 appears to have more occlusion than its neighboring fragments and recognize that the structure near the PSNP surface may look quite different from the native structure.

Although using methyl labeling is a versatile technique, it clearly has challenges for characterizing the dynamic binding of proteins to nanoparticles. Methyl labeling of lysine residues is rapid, which can be advantageous where a fast modification is desirable.^55^ However, one disadvantage is the fact degree of methylation on a single lysine cannot be easily controlled. Even with a large stoichiometric excess, some lysine residues may only be monomethylated during the short reaction time used. Acetylation could potentially be beneficial in these circumstances since this reaction can occur only once per lysine.^56^ However, the acetyl group is larger than the methyl group and, as a result, has a more significant steric hindrance. Because of this steric hindrance, the acetyl group might not be able to access lysine residues that are only partially exposed or are somewhat buried within the protein’s structure. Acetylation also neutralizes the lysine’s charge, which may lead to altered nanoparticle binding affinity and dynamics. Moreover, acetylation will not solve the key structural challenge described above of identifying whether a protein remains folded when adsorbed. In this sense, it suffers from the same drawback that methylation does.

### Altered Hydrogen Exchange Rates Identify Both Stabilized and Destabilized Regions in R2ab

Because of the challenges involved with methylation, we next sought to determine whether NMR-based hydrogen-deuterium exchange (HDX) could identify structural differences in the presence of PSNPs. Real-time HDX offers several potential advantages over methylation; namely, there is no direct modification of the protein, and the exchange can be monitored in real-time. HDX has been used before to monitor protein properties in the presence of PSNPs,^57,58^ and it was shown that the presence of gold nanoparticles made no measurable difference on the HDX rates for two model proteins, Ubiquitin and GB3.^28^

HDX-NMR is a powerful technique used primarily to study the dynamics, interactions, and conformational changes of proteins and other biomolecules. It relies on the principle that labile hydrogen atoms in proteins can be exchanged with deuterium atoms when exposed to a deuterated solvent, typically D_2_O.^59–61^ The rate at which this exchange occurs (k_ex_) can be indicative of the local environment surrounding specific amino acid residues. When proteins are folded in their native state, buried backbone protons can be protected from HDX. If the secondary structure is disrupted or if the protein topology is perturbed, the HDX rates will change.

R2ab has a total of 156 residues, and it was observed that many of the residues decayed very quickly. Two experimental considerations also reduced the number of observable number of observable independent spin systems. First, low concentrations (100 μM after adding D_2_O) needed to be used to prevent aggregation of R2ab in the presence of NPs; this increased the signal averaging required for each SOFAST-HMQC spectrum. Second, in order to collect spectra quickly, a small number of indirect time points were used. The resulting experiments (16 scans per increment, 128 total increments) represent a compromise between speed and sensitivity, with each HMQC requiring approximately 5 min to collect. Between these considerations and the intrinsically fast HDX observed in R2ab, we were only able to obtain measurements of k_ex_ both with and without PSNPs for only 35 residues of the protein. An additional 8 residues were only observable with PSNPs (Supporting Information, **Table S1**). This accounts for 22% of the protein. Alternatively, this value rises to 28% if the assumption is made that the 8 undetectable residues without PSNPs exchange faster than 0.05 min^-1^ (the fastest reliably detectable time in our experiments). While more complete coverage would be preferable, the detectable residues serve as probes of residue behavior throughout the protein structure.

All the HDX profiles for the selected residues were found to obey single exponential curves. The NMR signal intensity of the selected peaks was found to decrease relatively rapidly; however, because the final sample contains approximately 25% H_2_O, most signals did not decay entirely to zero. Examples of three such residues (**Figure 3**) demonstrate that exchange occurs rapidly (typically within 60-120 min) and all peaks have decayed to their baseline values by the end of the experiment. No exceptionally slow exchanging residues were observed, suggesting that, unlike established model proteins like cytochrome *c*,^62,63^ which can experience exchange times of days or longer, there are no highly protected residues in R2ab. Indeed, R2ab as a whole exchanges very quickly, and this may be influenced by its high lysine content (11%), which accelerates exchange.^64^

**Figure 3.**
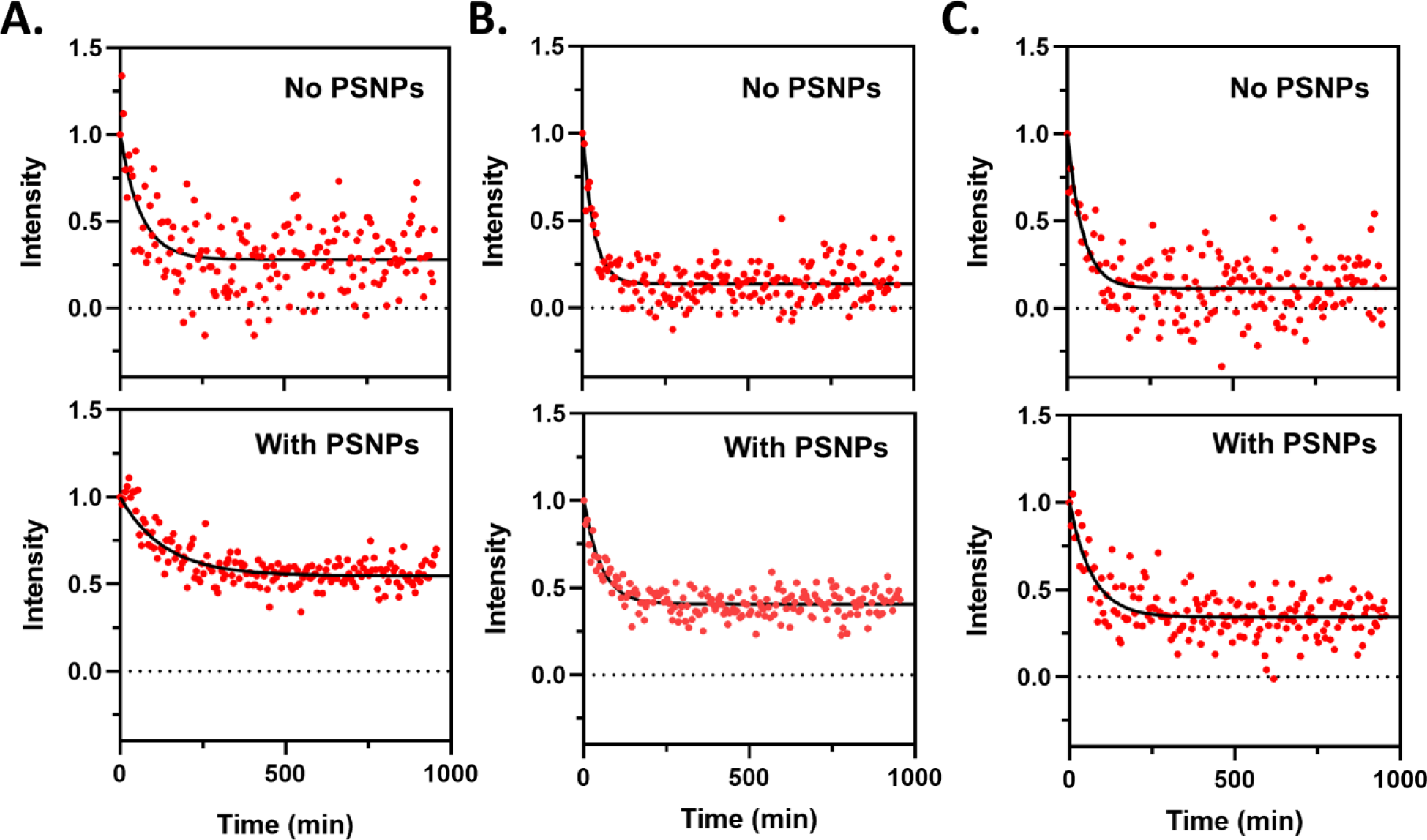
HDX kinetics of the residues 701 **(A)**, 785 **(B),** and 844 **(C)** in the absence (top) and presence (bottom) of PSNPs. Red points represent the measured intensity of each 2D NMR peak for each assigned residue after normalization of the initially collected SOFAST-HMQC time point. The black line represents the fit of the exponential decay as described in the text.

While there are some exceptions, most of the observed residues exhibit slower exchange in the presence of PSNPs (**Figure 4A-B**). This is somewhat surprising given that R2ab is expected to be destabilized on the nanoparticle surface;^8^ however, this effect is consistent with a reduction in methylation observed in many of the fragments (e.g., 699-717 and 771-783; **Figure 1A**). The rank order of residues is largely retained, and a correlation between HDX rates with/without PSNPs shows a linear trend for most residues (**Figure 4C**). This may reflect regions of partial stability in the adsorbed state, where the native structure can be retained. The average ratio of k_ex_ with/without PSNPs is 0.68, and many residues fall within a standard deviation of this average. Several notable exceptions to this trend occur, however. Residues 744, and 747, and Q802 exhibit the largest increase in k_ex_ relative to average behavior when PSNPs are present. Residue 802 sits at the interface between the R2a and R2b subdomains, and it destabilization of this interface is reasonable if the protein fold is disrupted. Residues L744 and K747 exist in a tight turn near the end of the R2a subdomain. Interestingly both of these regions are spatially surrounded by some of the slower exchanging residues; residue 802 is surrounded by H793, a residue that experiences significantly slower exchange in the presence of PSNPs. Residues 744 and 747 are flanked by T720, Y722, and T743, which also have slower exchange when PSNPs are added. K787, a residue with a large decrease in the presence of PSNPs, is located near N785 and T846, residues with a slight increase relative to the average. Overall, there are no obvious patterns; e.g., the R2a subdomain does not have an obviously slower k_ex_ in the presence of PSNPs relative to R2b. Instead, there are clusters of more quickly exchanging residues surrounded by residues with a slower-than-average k_ex_ rates(**Figure 4D**).

**Figure 4.**
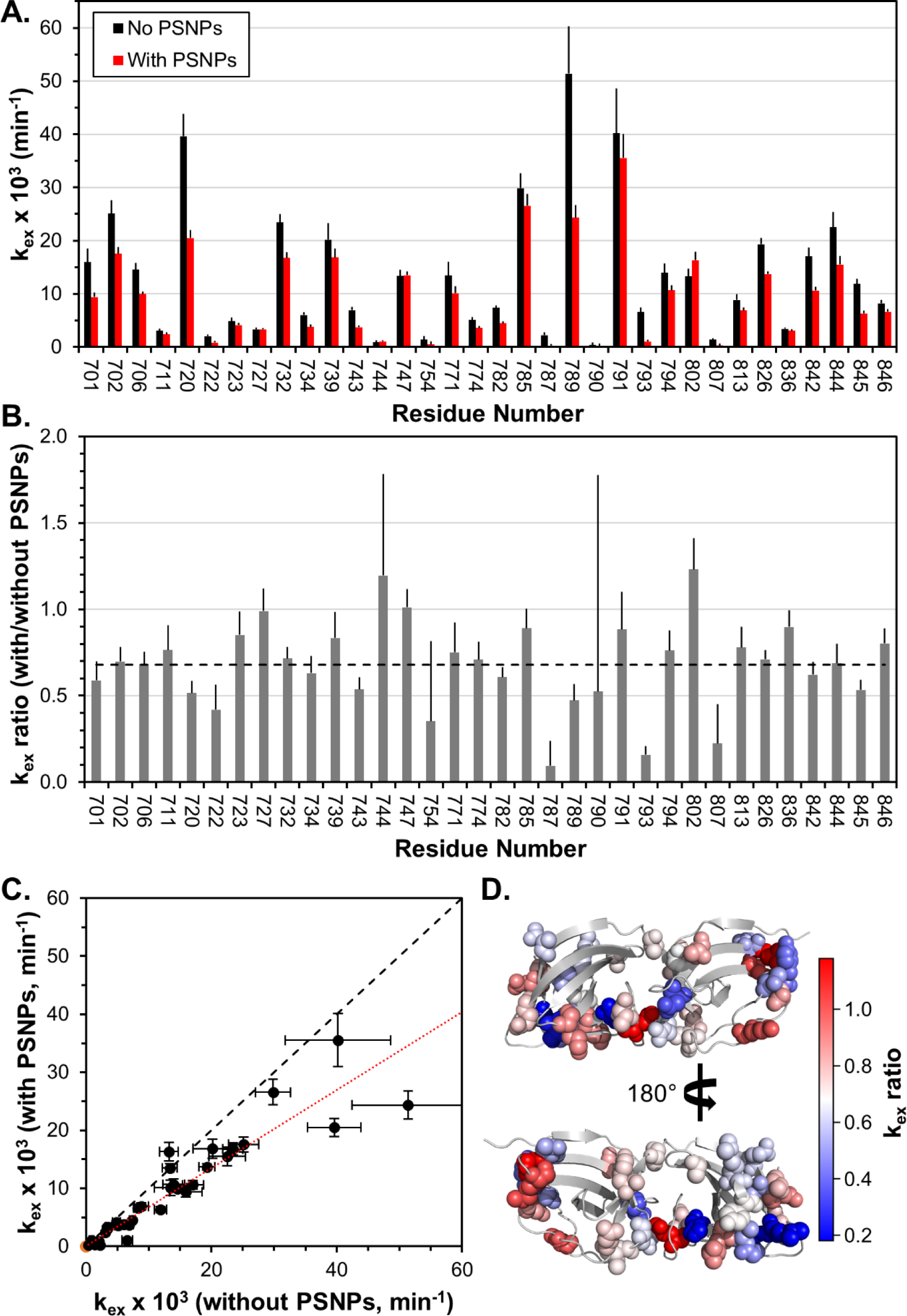
**(A)** HDX rates (k_ex_) for R2ab where rates could be compared with (red) and without (black) PSNPs. **(B)** Ratios of k_ex_ values for each residue; larger rates represent faster exchange in the presence of PSNPs. A black dashed line represents the average rate. **(C)** Correlation between rates taken from (A). The black dashed line represents the equation *y* = *x*. The dotted line represents the best fit of rates through the origin. **(D)** A comparison of rates mapped onto the structure of R2ab. Detectable residues are shown as CPK spheres. Residues with faster HDX rates relative to the average k_ex_ ratio are represented on the R2ab structure in red, while those with slower rates are shown in blue. Neutral residues, similar to the average shown in (B), are plotted in white.

Given the sparseness of the data, it is not surprising that most of the residues exchange more slowly: Given that many of the residues in the absence of PSNPs already exchange fairly quickly, detecting residues where k_ex_ increases would be difficult. Nevertheless, the clustering of slower/faster residues is suggestive. In the adsorbotope model, a handful of specific (and possibly exchanging) unfolded anchor points tether the protein to the nanoparticle surfaces. These anchor points are separated by sections of structures that may exhibit partial stability. The overall decrease in HDX rates can be explained if the anchors and general steric occlusion near the surface reduce proteins ability to adopt extended disordered conformations. However, regions far from anchor points could still experience faster than-average k_ex_ rates. Moreover, the clustering of faster/slower regions may reflect the underlying structure of the anchors. Mutagenesis may help reveal the physical characteristics of these adsorbotope anchoring sites. Overall, it can be suggested that the adsorption of R2ab onto PSNPs may give rise to an unfolding intermediate, with local protection in regions between the adsorbotope anchors. A localized protection effect of this nature has been observed before by Engel *et al.* for bovine α-lactalbumin on PSNPs.^65^

As with the methylation data, the large uncertainties and the subtle differences observed in HDX data emphasize the challenges in the structural interpretation of biophysical data for dynamically adsorbed proteins on nanoparticles. Only a handful of HDX rates can be reliably measured, and the heterogeneity in the adsorbed ensemble makes interpretations of the comparisons challenging. Prior to this, Sheibani *et al.* attempted to characterize the corona of gold nanoparticles using cryo-electron microscopy.^66^ They observed a density consistent with an ensemble of protein orientations and conformations. While the approaches employed here directly sample the solution state of nanoparticle binding *in situ*, both methylation and HDX rates reveal a similar picture: Experimental observations are varied, optimizing conditions that facilitate measurement is challenging, and clear trends are often lacking. Despite this, understanding protein-surface interactions remains a high priority. Methylation frequencies and HDX rates are useful metrics that sample protein behavior at the level of individual residues. Combining these with other tools such as NMR relaxation^67^, FPOP^50,51^,and proteolysis,^5,49^ may yet reveal the weak conformational determinants that underlie protein-surface interactions.

## Conclusions

Understanding the structural consequences of protein-surface interactions is important for understanding the nanoparticle corona, and it is also highly relevant for other fields, such as the initial attachment in biofilm formation. Here, we have examined the properties of R2ab, a domain implicated in bacterial attachment to polystyrene surfaces and known to interact strongly with PSNPs. To extend the capabilities of ensemble-based measurements like CD and ITC, we focused on approaches that provide structural information at the level of individual residues. Two approaches were employed: In the first, we performed compared lysine methylation rates in the presence and absence of PSNPs, quantifying methylation using LC-MS. In the second, we compared HDX rates with and without PSNPs using real-time NMR spectroscopy. Both techniques used here provide unique perspectives on protein structure and orientation, but they also present unique challenges, especially for a protein like R2ab that tends to induce aggregation in protein-nanoparticle mixtures. Methylation of R2ab occurs very quickly and obtaining large differences in partial methylation rates is a challenge. These reactions yield small differences that are interpretable but less than ideal. In HDX-NMR, fast exchanging residues and instrument sensitivity limit the number of residues that can be directly compared. In a dynamic environment such as R2ab interacting with PSNPs, a majority of data points go undetected, and those that are detected may reflect an ensemble average of several populations. Despite these challenges, we find that methylation and HDX produce useful information about the R2ab-PSNP interaction: Methylation identifies several fragments that exhibit small but statistically significant protection in the presence of PSNPs, presumably because of steric occlusion. HDX rates are observed to both increase and decrease in the presence of AuNPs. In light of our recently proposed adsorbotope model, where disordered anchor sites are interspersed with partially unfolded intervening regions, we expect that regions near anchor sites would exhibit accelerated HDX, whereas regions intervening regions may be somewhat stabilized by confinement near a surface. Overall, when combined with other structural approaches, HDX, and lysine methylation can complement studies of protein structure in the nanoparticle corona.

## Author Contributions

R.P.S., J.J.K. and N.C.F. conceived the idea and designed the experiments. R.P.S., S.K.M., J.J.K., J.S.S., and N.C.F. developed the methodology. R.P.S., S.K.M., C.S.K. and N.C.F. performed the experiments, R.P.S., S.K.M., J.S.S., and N.C.F. analyzed the results. R.P.S. and N.C.F. wrote the original manuscript. R.P.S., S.K.M., C.S.K., J.J.K., J.S.S., and N.C.F. revised and edited the manuscript. All authors approved the final version of the manuscript.

## Supporting information

All Supporting Information

## Acknowledgments

We thank Tim Dowell for a helpful presentation on crafting scientific illustrations. This work was supported by the National Institute of Allergy and Infectious Diseases of the National Institutes of Health under grant numbers R01AI139479 (to NCF), and the National Science Foundation under grant number MCB 1818090 (to NCF). JSS and SKM acknowledge support from the Glycoscience Center of Research Excellence (NIH P20GM103460).

