## Supplementary material for "Exploring the Residue-Level Interactions between the R2ab Protein and Polystyrene Nanoparticles": All Supporting Information

**Supporting Table**

**Table S1.** HDX Exchange Rates for R2ab with and without PSNPs.

Rates that could not be measured reliably are provided with a “-“. These rates are likely greater than 0.05 $min^{-1}$, the fastest $k_{ex}$ rate reliably measurable in our approach.

| Residue | Amino Acid | No PSNPs $k_{ex}$ x $10^3$ ($min^{-1}$) | With PSNPs $k_{ex}$ x $10^3$ ($min^{-1}$) |
|---|---|---|---|
| 701 | V | 16 ± 3 | 9.4 ± 0.9 |
| 702 | T | 25 ± 2 | 17.6 ± 1.3 |
| 706 | G | 14.5 ± 1.3 | 9.9 ± 0.5 |
| 710 | I | - | 0.2 ± 1.0 |
| 711 | N | 3.1 ± 0.4 | 2.4 ± 0.3 |
| 716 | G | - | 0.5 ± 0.3 |
| 720 | T | 40 ± 4 | 20.5 ± 1.5 |
| 722 | Y | 2.0 ± 0.4 | 0.9 ± 0.3 |
| 723 | D | 4.9 ± 0.7 | 4.1 ± 0.4 |
| 727 | K | 3.3 ± 0.3 | 3.3 ± 0.3 |
| 732 | I | 23.4 ± 1.6 | 16.8 ± 1.1 |
| 734 | R | 6.0 ± 0.5 | 3.8 ± 0.5 |
| 737 | S | - | 2.6 ± 0.5 |
| 739 | T | 20 ± 3 | 16.8 ± 1.7 |
| 743 | T | 6.9 ± 0.6 | 3.7 ± 0.4 |
| 744 | L | 0.9 ± 0.4 | 1.0 ± 0.2 |
| 747 | K | 13.4 ± 1.2 | 13.5 ± 0.8 |
| 754 | D | 1.4 ± 0.7 | 0.5 ± 0.6 |
| 761 | Y | - | 0.5 ± 0.6 |
| 762 | G | - | 110 ± 20 |
| 771 | Y | 13 ± 3 | 10.1 ± 1.3 |
| 772 | Q | - | 16 ± 2 |

| Residue | Amino Acid | No PSNPs $k_{ex}$ x $10^3$ ($min^{-1}$) | With PSNPs $k_{ex}$ x $10^3$ ($min^{-1}$) |
|---|---|---|---|
| 773 | T | - | 59 ± 8 |
| 774 | A | 5.1 ± 0.5 | 3.6 ± 0.4 |
| 782 | Q | 7.4 ± 0.4 | 4.5 ± 0.3 |
| 785 | N | 30 ± 3 | 27 ± 2 |
| 787 | K | 2.2 ± 0.6 | 0.2 ± 0.3 |
| 789 | G | 51 ± 9 | 24 ± 2 |
| 790 | V | 0.4 ± 0.4 | 0.2 ± 0.4 |
| 791 | K | 40 ± 8 | 36 ± 5 |
| 793 | H | 6.6 ± 0.9 | 1.0 ± 0.3 |
| 794 | T | 14.0 ± 1.7 | 10.6 ± 1.0 |
| 802 | Q | 13.3 ± 1.5 | 16.3 ± 1.6 |
| 804 | A | - | 0.3 ± 0.4 |
| 807 | V | 1.4 ± 0.3 | 0.3 ± 0.3 |
| 813 | Q | 8.8 ± 1.1 | 6.9 ± 0.6 |
| 826 | A | 19.3 ± 1.3 | 13.7 ± 0.5 |
| 829 | L | - | 0.2 ± 0.7 |
| 836 | K | 3.4 ± 0.3 | 3.1 ± 0.2 |
| 842 | K | 17.0 ± 1.7 | 10.6 ± 0.7 |
| 844 | Y | 23 ± 3 | 15.5 ± 1.6 |
| 845 | L | 11.8 ± 1.0 | 6.3 ± 0.5 |
| 846 | T | 8.2 ± 0.6 | 6.6 ± 0.5 |

**Supporting Figure**

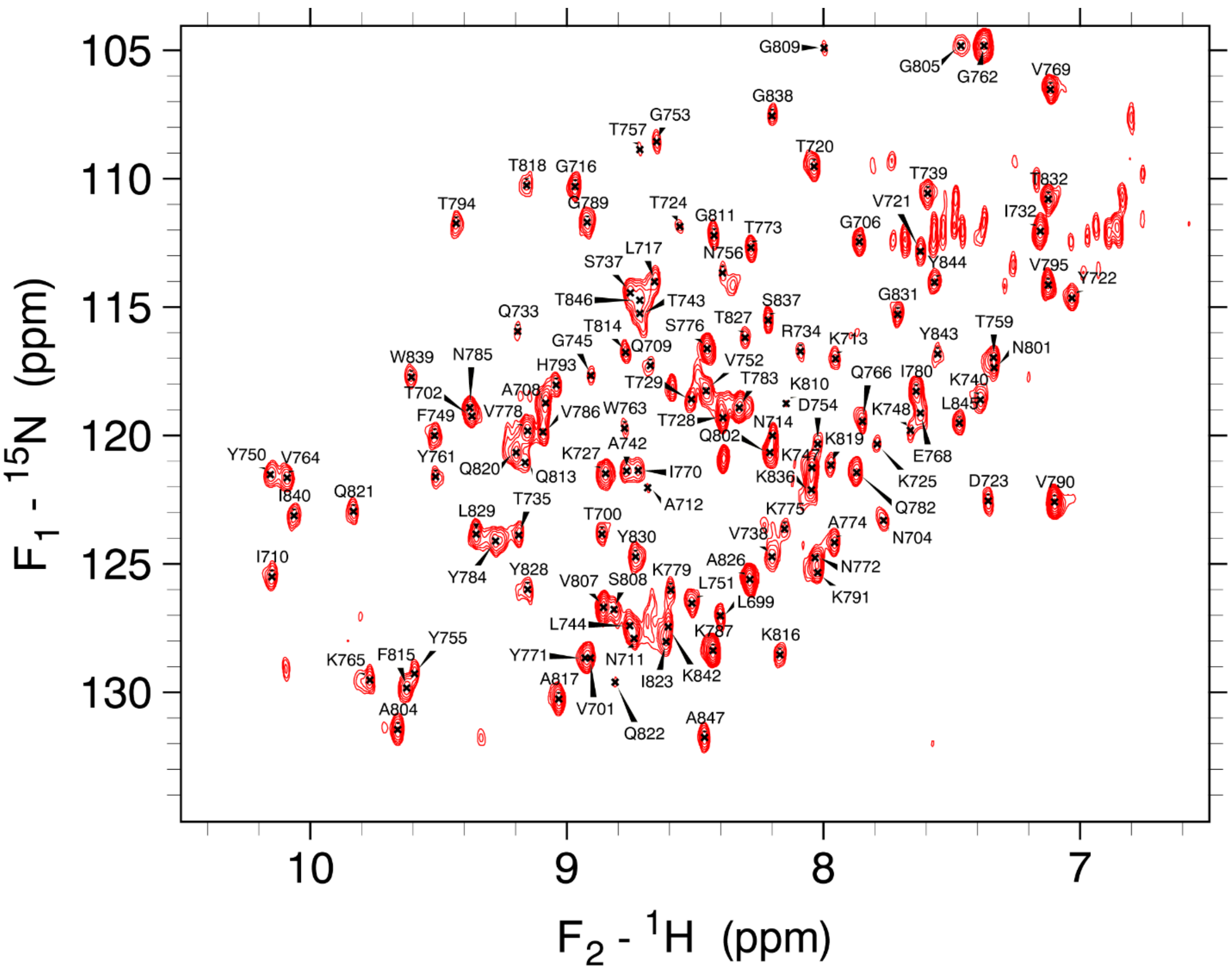

**Figure S1.** NMR peak assignments for R2ab at pH 4.5.

SOFAST-HMQC spectrum of 100 μM $^{15}$N-labeled R2ab. The peak assignments were transferred from a spectrum collected at pH 6.5 (BMRB ID 30821).
